# The evolutionarily conserved C-terminal domain of a domesticated transposase-derived protein regulates its DNA integration ability

**DOI:** 10.64898/2026.08.31.747927

**Authors:** Aditi Saha, Aryaman Ghosh, Sharmistha Majumdar

## Abstract

THAP9 is a transposable element-derived gene which encodes a protein that is homologous to the active *Drosophila* P-element transposase (DmTNP). Both THAP9 and DmTNP possess a C-terminal domain (CTD) which is functionally uncharacterized.

Sequence and structural analysis suggest that the THAP9-CTD has a novel fold which is only found in THAP9 homologs. To explore the evolutionary history and characteristics of this novel domain, exhaustive phylogenetic analysis (using MSA, structure prediction, MSTA-based clustering) was performed. THAP9-CTD homologs were more widely distributed throughout the animal kingdom in comparison to DmTNP-CTD homologs which were restricted to arthropods. Moreover, the THAP9-CTD homologs were more conserved, especially among mammals and birds and their average length increased in a class-specific manner. Comparison with the DmTNP-CTD homologs demonstrates that although their respective CTDs may have evolved independently, they both surprisingly share similar secondary structure elements consisting of three conserved helical regions made of hydrophobic residues that are predicted to make up a conserved core.

The role of the respective CTDs were further investigated by creating truncation mutants lacking the CTD. Interestingly both THAP9 and DmTNP truncation mutants are still capable of DNA excision and integration, suggesting that their respective CTDs are not essential for DNA transposition. Moreover, CTD truncation favours DNA integration in THAP9: this suggests that CTD acquisition during evolution may have led to THAP9’s domestication as observed in other transposable element-derived genes like Rag1 and piggybac, which have similar terminal regulatory domains.

## Introduction

Transposable elements or mobile genetic elements are segments of DNA which can move from one position to another within a genome. This movement known as transposition can be both beneficial and detrimental for the organism. Over the course of evolution, some transposons continue to actively mobilise while others lose their mobility by becoming stationary within a genome and may acquire new functions. Human THAP9 is one such transposase-derived gene which is speculated to have lost its ability to jump. The encoded protein closely resembles the active *Drosophila* P-element transposase (DmTNP), bearing 25% sequence identity and 40% homology.

Human THAP9 (hTHAP9) is a transposase-derived gene, encoding a protein which recognizes the terminal inverted repeats (TIRs) of DmTNP and mobilizes in *Drosophila* as well as HEK293 cells *in vivo* (Majumdar et al. 2013). The hTHAP9 protein has 4 domains: N-terminal THAP domain (NTD), oligomerization domain, RNaseH like catalytic domain, and C-terminal domain (CTD). Although it is possibly domesticated i.e. immobile in the human genome, THAP9 retains its catalytic ability to excise and integrate the P-element transposon.

THAP9 is a member of the THAP family of proteins which all share the characteristic THAP domain that can bind DNA via a signature C2CH zinc coordinating motif. However, THAP proteins have diverse cellular functions. Many THAP proteins have additional domains. For instance, the HCF1 binding motifs in THAP1-7,9,11, nitrobindin domain in THAP4 and RNaseH-like domain in THAP9. Moreover, some THAP proteins like DmTNP and THAP9 and THAP11 (as per InterPro) have uncharacterized carboxy-terminal domains (CTD). These CTDs do not appear to be related to each other and have low sequence conservation as shown by their pairwise alignment (Fig.1).

**Figure 1:**
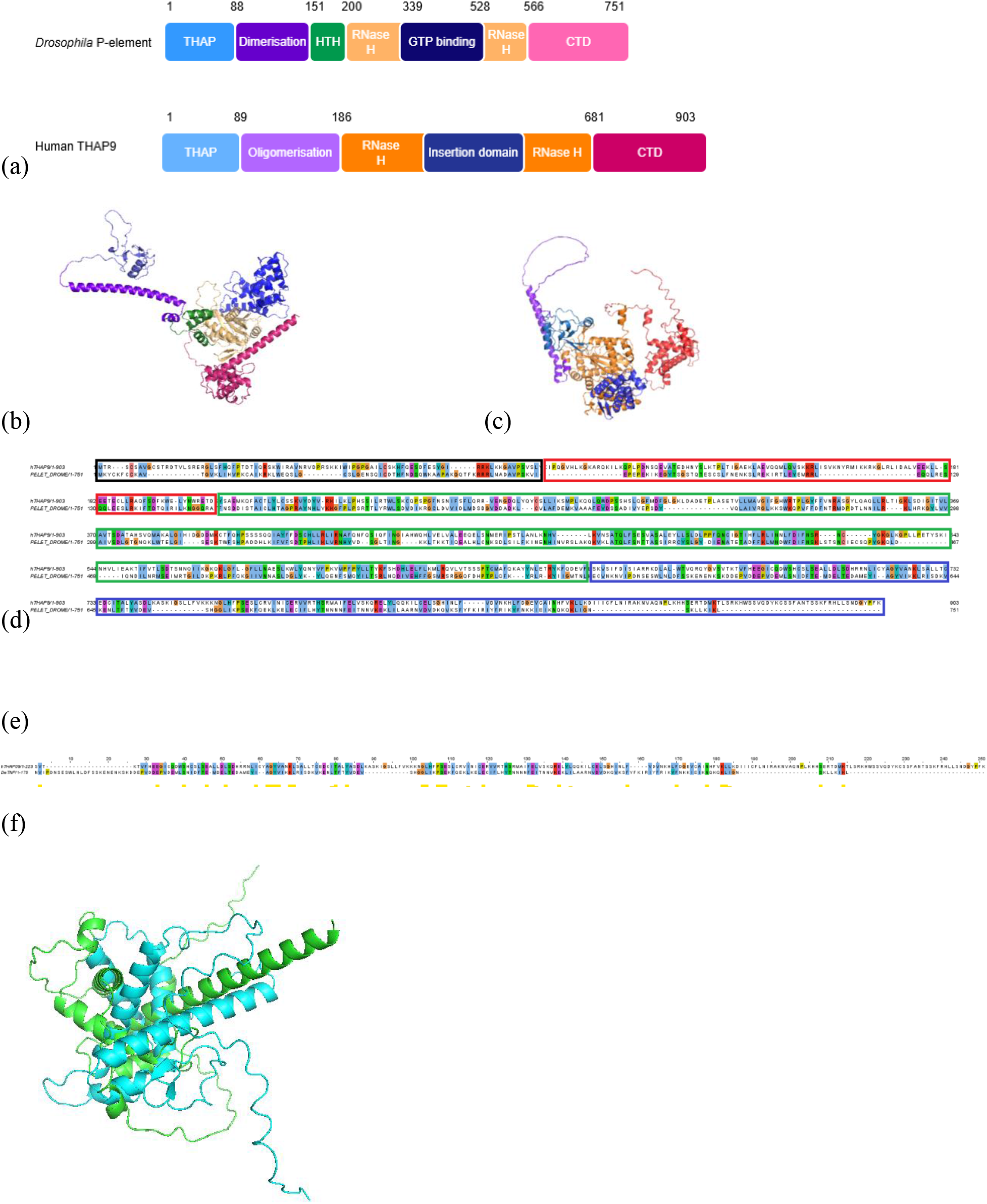
(a) Schematic representation of domain architecture with domain boundaries. Alphafold predicted structures of (b) DmTNP and (c) hTHAP9. The colour coding in (a) corresponds to the respective domains in (b) and (c). (d) Pairwise alignment of hTHAP9 and DmTNP highlighting the oligomerisation domain (red box), RNaseH like domain (green box), THAP domain (black box) and CTD (blue box). (e).Pairwise sequence alignment of C-terminal domains (from InterPro) of hTHAP9, DmTNP (f) Structural superposition of the predicted CTDs of DmTNP (green) and hTHAP9 (cyan).

Most transposases are multi-domain proteins. In addition to specific domains involved in DNA binding, cleavage and integration, some transposases have carboxy-terminal domains which have diverse regulatory roles. For example, the Rag1 domesticated transposase, which is involved in V-(D)-J recombination, has a C-terminal domain which plays an autoinhibitory role during cleavage of single stranded DNA but not double stranded DNA (De et al. 2004). Interestingly, the active protoRag transposase (a primitive version of Rag1, present in lower organisms) possesses a C-terminal tail (in addition to a C-terminal domain) which is involved in DNA cleavage and loss of which contributes to the domestication of Rag1 (Tao et al. 2020; Zhang et al. 2019). Experimental reconstructions demonstrate that inclusion of the ancestral CTD restores robust transposition activity, whereas its absence in vertebrate RAG1 abolishes transposition but preserves precise DNA cleavage at RSSs. On the other hand, the cysteine rich C-terminal domain of PiggyBac transposase, is not necessary for transposition, but triggers and fine tunes the preciseness of DNA cleavage as well as integration (Helou et al. 2021). However, the exact role of the C-terminal domain of THAP9 is not yet known.

Here we investigate the C-terminal domain of hTHAP9 and DmTNP and characterise their role in the excision and integration of the *Drosophila* P-element DNA. We also carried out a detailed evolutionary analysis of the CTDs to determine the extent of its conservation and divergence across different taxonomic groups as well as to identify conserved residues and structural elements.

## Materials and Methods

### Data Collection

The CTD of hTHAP9 was named “DNA transposase THAP9, C-terminal domain” (IPR055035) in the InterPro database (Blum et al. 2025) and extended from residues 681-903. Interpro listed 647 entries (corresponding to THAP9-CTD homologs) with only one reviewed entry (Q9H5L6, THAP9 *Homo sapiens*). On the other hand, InterPro listed the CTD of DmTNP as “Transposable element P transposase-like, C-terminal” (IPR022242), which corresponded to residues 567-751 and had 709 entries with only one reviewed entry (Q7M3K2, Transposable Element P transposase, *Drosophila melanogaster*). In this study, the homologs of DmTNP-CTD were termed as TNP-CTD and the homologs of hTHAP9-CTD were labelled as THAP9-CTD.

To obtain the sequences for individual CTD homologs, search was performed using phmmer (Rajković et al. 2026). The retrieved sequences were obtained, keeping the database as UniProt, and e-values and Hit-Scores as 1. Multiple sequence alignment with the obtained sequences served as input to hmmbuild. The resultant hmmprofile was then used as a query to identify additional CTD domain sequences in Uniprot. The sequences were downloaded from Uniprot, trimmed as per the predicted domain boundaries and JSON files (containing metadata) for the resultant data were retrieved from Uniprot. Next, Whole Genome Shotgun (WGS) sequenced assemblies were removed from the InterPro datasets. WGS are sequences, which are draft contigs generated by shotgun sequencing and subsequent de-novo assembly, may be of low-quality, contaminated, redundant or inaccurate, and removing them makes the dataset more robust and consistent. After removing WGS sequences, the dataset was reduced to 623 for THAP9-CTD and 267 entries for TNP-CTD.

### Data organisation

The collected dataset (FASTA file) was arranged as per taxonomic hierarchy. The metadata for taxonomic details of each dataset was obtained from the NCBI Taxonomy Database (https://eutils.ncbi.nlm.nih.gov/entrez/eutils/efetch.fcgi?db=taxonomy&id=tax_id&retmode=xml); (O’Leary et al. 2024). The dataset was arranged in the order – “Kingdom”, “Phylum”, “Class”, “Order”, “Family” and “Genus”. CD-HIT (Fu et al. 2012) was performed on the dataset at 80% similarity to remove redundant entries. For this, the FASTA files of each dataset were disassembled for each individual species (using a custom python script) and after performing CD-HIT for all the species individually, all the sequences were reassembled. The Uniprot database removed several proteins from their database in February 2026, and hence the number of sequences further reduced to 39 for DmTNP-CTD and 173 for THAP9-CTD.

### Data visualisation

The MAFFT algorithm (Katoh and Standley 2013) was used for creating Multiple Sequence Alignment (MSAs) with the FASTA files, to identify conserved regions as well as possible class-specific mutations. Jalview (Waterhouse et al. 2009) was used for visualization of the MSA, with a colour scheme based on the nature of amino acid as per the Clustal colours.

Clustal X Default Colouring
| Category | Colour | Residue at position | { Threshold, Residue group } |
| --- | --- | --- | --- |
| Hydrophobic | BLUE | A,C,I,L,M,F,W,V | {>60%, WLVIMAFCYHP} |
|  |  | C | {>60%, WLVIMAFCYHP} |
| Positive charge | RED | K,R | {>60%,KR},{>85%, K,R,Q} |
| Negative charge | MAGENTA | E | {>60%,KR},{>50%,QE},{>50%,ED},{>85%,E,Q,D} |
|  |  | D | {>60%,KR},{>85%, D,E,N},{>50%,ED} |
| Polar | GREEN | N | {>50%, N},{>85%, N,D} |
|  |  | Q | {>60%,KR},{>50%,QE},{>85%,Q,T,K,R} |
|  |  | S,T | {>60%, WLVIMAFCYHP},{>50%, TS},{>85%,S,T} |
| Cysteines | PINK | C | {>85%, C} |
| Glycines | ORANGE | G | {>0%, G} |
| Prolines | YELLOW | P | {>0%, P} |
| Aromatic | CYAN | H,Y | {>60%, WLVIMAFCYHP},{>85%, W,Y,A,C,P,Q,F,H,I,L,M,V} |
| Unconserved | WHITE | any / gap | If none of the above criteria are met |

Using the MSA created by MAFFT, phylogenetic trees were created using IQ Tree (Nguyen et al. 2015) which uses Maximum Likelihood. The parameters were kept as default and 1000 bootstraps were performed for both the datasets. The resultant tree was then visualized, coloured and analyzed using the itol online tool.

### Sequence clustering

After performing CD-HIT, clustering of data was done using DIAMOND at 40% similarity threshold to get representative sequences based on similarity (Buchfink et al. 2026). Therefore, two rounds of clustering had been performed: first based on species using CD-HIT at 80% similarity, and the second clustering was done based only on sequence using DIAMOND at 40% similarity. The resultant TNP-CTD dataset had 38 clusters while the THAP9-CTD dataset had 75 clusters.

### Structure based alignment

Multiple Structural Alignment (MSTA) was performed to help find distant homologs between the clusters that may not have been observable by just sequence-based alignments. First using the centroid of each cluster, AlphaFold structures of the domains were generated using each centroid as the input sequence for structure generation in the AlphaFold server (Abramson et al. 2024). For clusters of each dataset, MSTA was performed using MUSTANG (Konagurthu et al. 2006). Using MSTA, RMSD (Root-Mean-square-Deviation) data was created and converted to a heatmap in Matplotlib using a custom python code (The Matplotlib Development Team 2026).

### Prediction for disordered regions

The intrinsic disorderliness of the domains was predicted using IUPred (Dosztanyi et al. 2005). The MSAs of CTDs of DmTNP and THAP9 as well as their homologs were uploaded in the webserver. The residue wise disorder scores were predicted via default parameter and plotted using custom python codes.

### Site directed Mutagenesis

Primers were designed using NEB Base changer tool^TM^ for generating THAP9 mutants in which either one or both terminal domains were truncated. Site directed mutagenesis was performed using Phusion polymerase (F-530S, Thermoscientific) and KLD enzyme (M0554S, NEB) as per manufacturer’s instructions. All mutant constructs were verified by Sanger’s sequencing at CCAMP, India.

### In-vivo Excision assay

HEK293 cells (0.5 × 10^6^, grown in DMEM (SH30243.01, Cytiva), supplemented with 10% fetal bovine serum (under standard cell culture conditions) were plated per well of 6 well plate and transfected at 70-90% confluency using Turbofect ^TM^ (R0531, Thermoscientific) transfection reagent, as per manufacturer’s protocol. Briefly, 1ug pISP2/Km reporter plasmid (which has a 0.6kb P-element insertion, that disrupts the kanamycin resistance gene) and 1ug THAP9 (wild type or mutants separately, cloned in pcDNA3.1(+)) or pBlueScript empty vector (negative control) were transfected in triplicate. The cells were harvested 48 hours after transfection and resuspended in 1X PBS (TL1006, HIMEDIA). The plasmids from the cells were isolated using GeneJET plasmid isolation kit (K0691, ThermoScientific) as per manufacturer’s protocol (Sharma et al., 2021).

To determine if successful excision of the P-element DNA has occurred, the segment of the pISP2/Km reporter plasmid, which is flanked by the P-element TIRs, was amplified using Phusion polymerase (F-530S, ThermoScientific) at annealing temperature of 67-69 °C using Veriti Applied Biosystems Thermal cycler. The PCR products were resolved in 0.8% agarose gel and visualised by ethidium bromide staining.

Excision Forward Primer: 5’-GTTGTGTGGAATTGTGAGCGG-3’

Excision Reverse Primer: 5’-CCGGATCGGTCCTCACGATG-3’

pISP2 Amp Forward: 5’ - AACATTTCCGTGTCGCCCTTA-3’

pISP2 Amp Reverse: 5’ - AGTGAGGCACCTATCTCAGC-3’

If P-element excision has taken place, the expected size of the product of the PCR performed on the repaired pISP2/Km reporter plasmid (after excision) is ∼200bp. On the other hand, if P-element excision has not occurred, the expected size of the product of the PCR performed on the unexcised pISP2/Km reporter plasmid is ∼750 bp. As a positive control, the Ampicillin resistance gene (present in the reporter plasmid) was PCR-amplified to indicate that comparable amounts of plasmid DNA template were present in the PCR samples with a band at ∼850 bp.

### In-vivo Integration assay

Cg4-Neo reporter plasmid (carrying the G418 resistance gene, which is transcribed from the SV40 promoter and is flanked by the TIRs of the P-element) was co-transfected with THAP9 (wild type or mutants separately, cloned in pcDNA3.1(+)) or negative control (pBluescript empty vector) in HEK293 cells (0.4-0.5 × 10^6^ cells per well of a 6-well plate) at 70-90% confluency with Turbofect ^TM^ (R0531, Thermoscientific) transfection reagent, as per the instruction manual (Sharma et al., 2021). Briefly, a total of 2 ug of plasmid DNA was transfected per well: 50 ng of CgNeo4, 1 ug THAP9 (wild type or mutants separately) /negative control and 950ng pBluescript empty vector. After 48 hours, the cells from each well were harvested, seeded into 10 cm dishes and allowed to adhere for 24 hours at 37 °C in a CO_2_ incubator. After 24 hours, media supplemented with G418 (0.5 mg/mL, G0349, TCI) was added to the cells. and selected for 2-3 weeks. The G418-resistant colonies were fixed using methanol, stained with crystal violet, and counted. The relative integration activity was computed as a percentage of activity of each mutant with respect to wild type THAP9. The values are normalised with wild type THAP9.

Relative Integration Activity= (No. of G418 resistant colonies for mutants *100)/ (No. of G418 resistant colonies for wild type THAP9)

Significant values were determined using a one-way ANOVA method followed by Dunn Sidak test (Sharma et al. 2021).

### Protein Immunoblotting

The wild type and the mutant THAP9 proteins were tagged with a C-terminal HA tag. Extracts from transfected HEK293 cells (THAP9 and mutants) were lysed and run on a 10% denaturing SDS gel in 1X Tris-glycine-SDS running buffer. Precision Plus Protein Dual Color Standard (10 – 250kDa, 1610396, Biorad) was used as a molecular size marker. The samples were transferred to a PVDF membrane by electroblotting using a standard gel transfer system. The membrane was blocked using 5% skimmed milk in Tris-Buffered Saline with Tween-20 (TBST), to remove nonspecific binding. The membrane was washed with 1X TBST and incubated with anti HA primary antibody (1:2000 dilution, SAB1306169-400UL, Sigma) overnight at 4°C, followed by incubation with HRP coupled secondary antibody (1: 5000 dilution, 1706515, Biorad) for 2 hours at room temperature. Detection was performed using Clarity ECL Western Blotting substrates (1705061, Biorad) on a Biorad Gel Documentation System.

## Results

### The C-terminal domains of both hTHAP9 and DmTNP are uncharacterised

The hTHAP9 protein is homologous to the active *Drosophila* P-element transposase (DmTNP). Both DmTNP and hTHAP9 can bind the terminal inverted repeats of the P-element transposon (Majumdar et al. 2013; Sharma et al. 2021) via an RNase H like catalytic domain (Majumdar et al. 2013; Sharma et al. 2021). However, the properties of their C-terminal domain is unknown.

The domain organization of wild type DmTNP and hTHAP9 can be noted from their schematic representation (Fig. 1a) as well as their AlphaFold predicted structures (Fig 1b and c). Pairwise sequence alignment of full length DmTNP and hTHAP9 (Fig. 1d) as well as only the CTD (Fig. 1e) of both the proteins reveal that the full length proteins are 21% identical and 36% similar while the oligomerization domains and RNaseH-like domains have 20% identity and 37% similarity. On the other hand, the CTDs (Fig. 1e) are 15% identical and 29% similar.

To investigate if the CTDs of DmTNP and hTHAP9 had structural conservation, their structures were then predicted using AlphaFold. Comparison of the predicted CTD structures by superposition illustrated that the individual CTDs do not overlap with each other (RMSD ∼14Å) suggesting structural divergence (Fig 1f).

### Homologs of DmTNP-CTD mostly occur in Arthropods while homologs of THAP9-CTD are widely distributed from Arthropods to Chordates

To investigate the evolutionary conservation of the uncharacterised CTD of both DmTNP and hTHAP9, homologous sequences were downloaded from UniProt and arranged taxonomically as per the hierarchy of Kingdom, Phylum, Class, Order, Family and Genus. CD-HIT was performed to remove redundant entries followed by Multiple Sequence Alignment (using MAFFT). This extensive analysis led to the identification of 39 homologs of DmTNP-CTD (i.e. TNP-CTDs) and 173 homologs of hTHAP9-CTD (i.e. THAP9-CTDs). It was intriguing to note that the DmTNP-CTD was not retrieved as a homolog of hTHAP9-CTD and vice versa. This suggests that the CTDs of the two proteins are evolutionarily distinct and may have evolved independently.

The TNP-CTDs are restricted to kingdom Animalia (Fig 2), mostly occurring in invertebrates of phylum Cnidaria (Anthozoa) and Arthropoda (Insecta, Arachnida). Interestingly, most TNP-CTDs (67%) occur in Class Insecta, which is also the class to which the active transposase DmTNP belongs.

**Figure 2:**
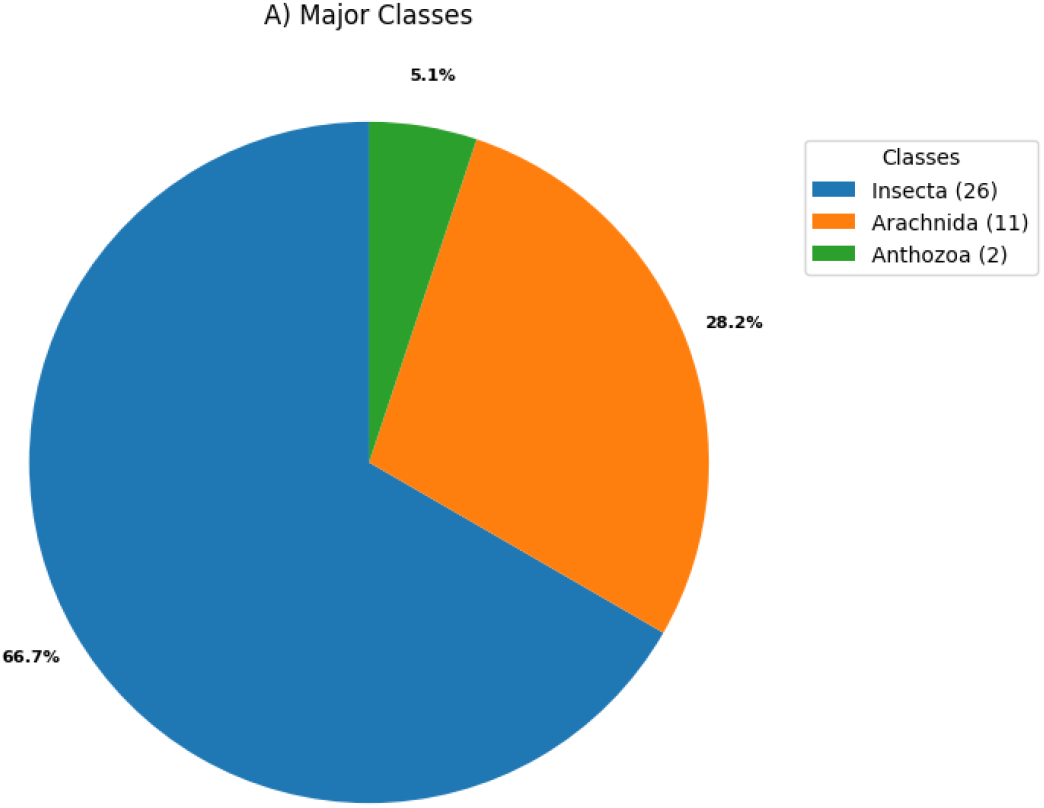
TNP-CTDs are restricted to Kingdom Animalia with maximum representation (67%) in Class Insecta

THAP9-CTDs are also restricted to Kingdom Animalia (Fig 3a); however these homologs are widely distributed across invertebrates ranging from phylum Arthopoda (Insecta, Arachnida, Malacostraca) to Chordata including vertebrates (Myxini, Actinopteri, Cladistia, Amphibia, Reptilia, Aves, Mammalia). Fig 3a represents the percentage of occurrence of THAP9-CTDs in the animal kingdom, with mammals possessing the maximum members (37.6%) within vertebrates while arachnids (24.3%) have the highest representation within invertebrates. Classes which have less representative members (crustaceans, hagfish, primitive bony fish) are clubbed as “Others”, details of which are illustrated in Fig 3b.

**Figure 3:**
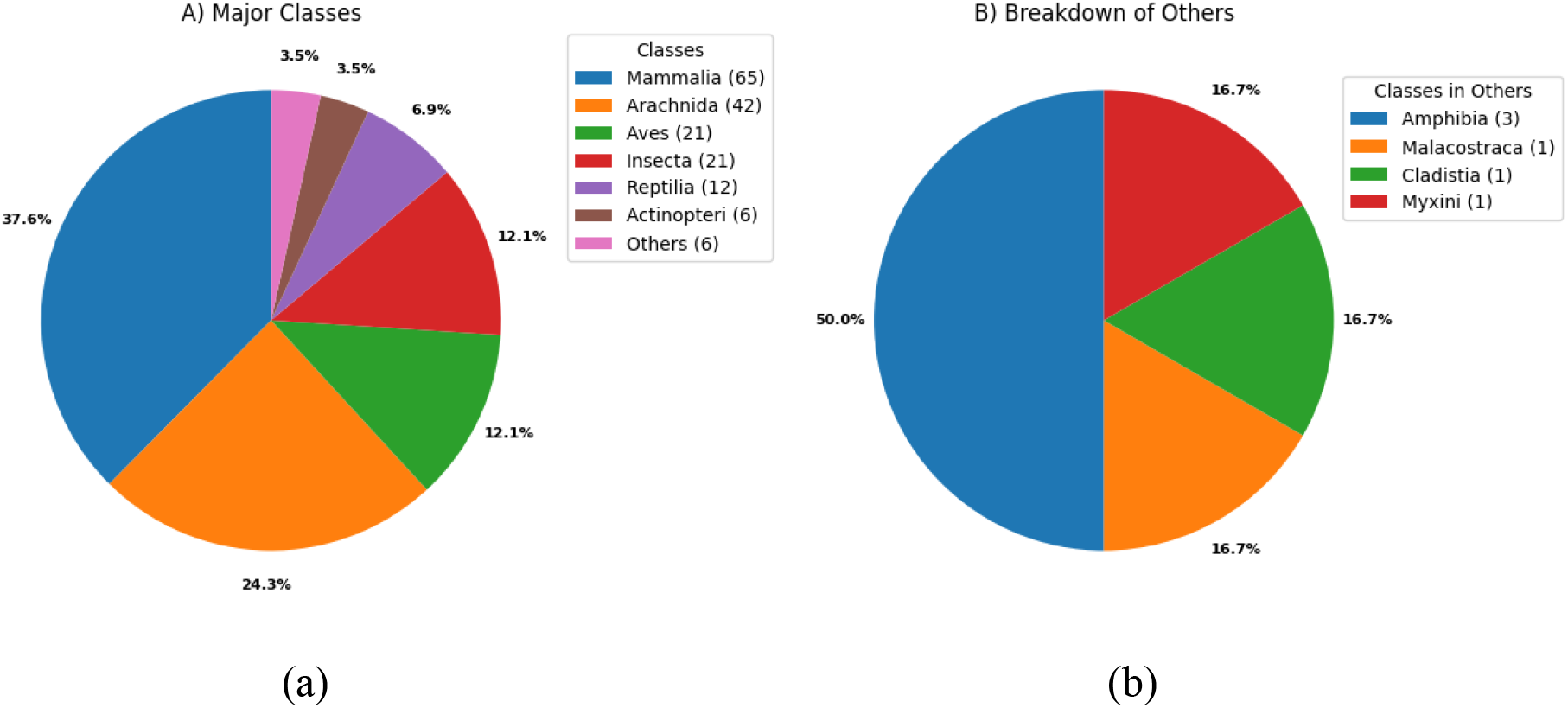
(a) THAP9-CTDs are spread across invertebrates and vertebrates with maximum representation in arachnids (24.3%) within invertebrates and Mammalia (37.6%) within vertebrates. (b) THAP9-CTDs are less represented in Crustaceans (Malacostraca), hagfish (Myxini) and primitive bony fish (Cladistia).

### Length distribution of THAP9-CTD and TNP-CTD

The lengths of CTDs (of THAP9 and TNP) were then investigated (Fig. 4). The Tropical sea anemone (A0A913X3G4) belonging to Anthozoa has the longest TNP-CTD (211 residues) whereas *Mustard beetle* (A0A9P0DWI1) from Insecta has the shortest TNP-CTD (60 residues). On the other hand, the average length of THAP9-CTDs are longest in mammals, which also have the highest representation (Fig. 4); however, the longest THAP9-CTD (233 residues) occurred in Three-toed box turtle (A0A674I1E7) from Reptilia while the shortest THAP9-CTD (81 residues) occurred in *Hirondellea gigas* (A0A6A7G904) from Malacostraca.

**Figure 4:**
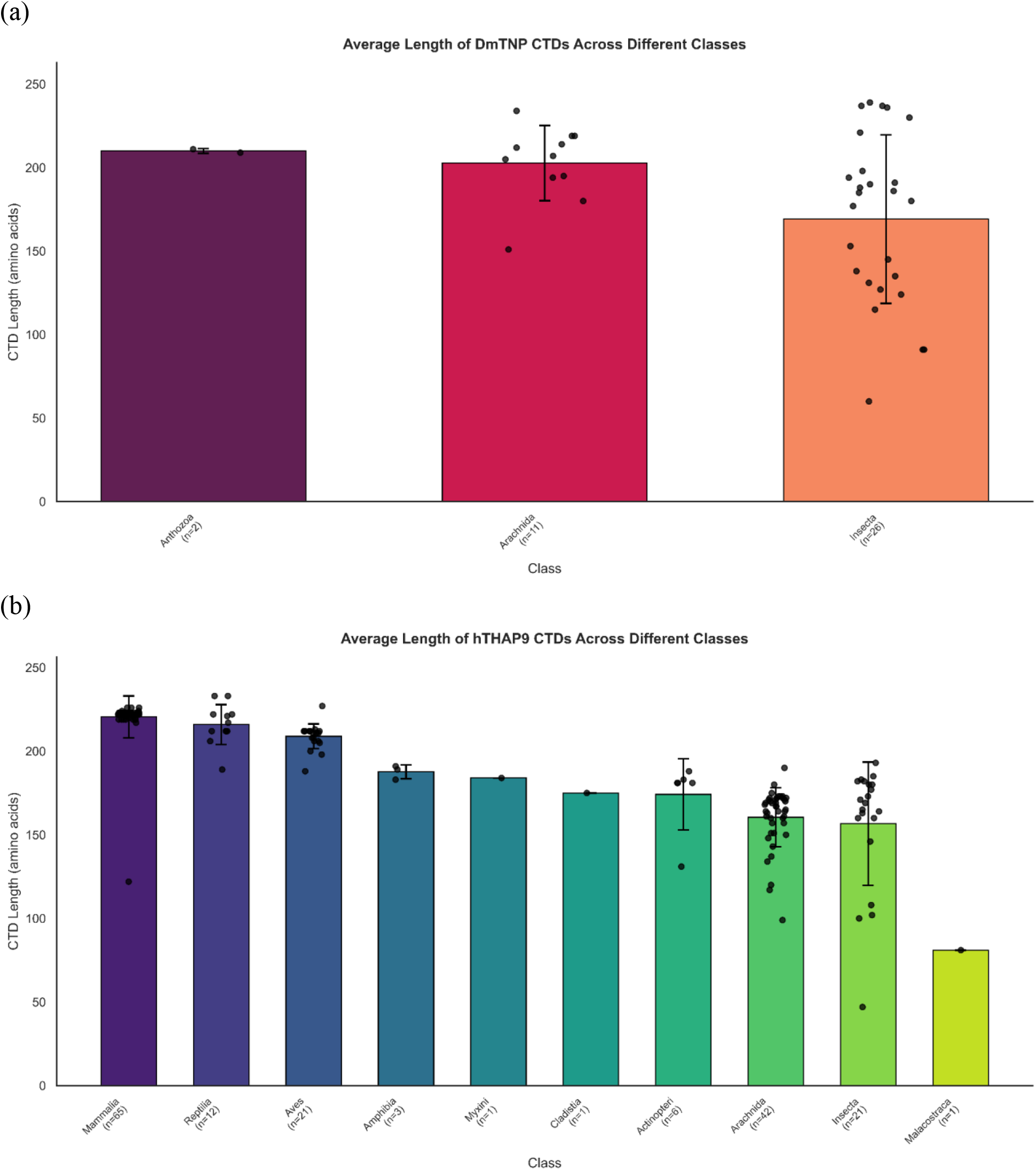
Average length of (a) TNP-CTDs and (b) THAP9-CTDs across classes where Anthozoa have the highest average length for TNP-CTDs and Mammalia have highest average length for THAP9-CTDs. Individual data points corresponding to lengths of individual CTDs are overlaid on the bars with the error bars representing standard deviation, X-axis representing the organism class and Y-axis representing the average length of CTDs.

Moreover, it is observed that the average length of THAP9-CTDs increases in a class-specific manner, indicating hierarchical increase of average CTD length during evolution. Similar class-specific trends are not observed for TNP-CTD since all members (except 2) belong to Arthropoda.

### Phylogenetic Analysis of the TNP-CTDs and THAP9-CTDs

To study the evolutionary history of both THAP9-CTD and TNP-CTD, phylogenetic trees were constructed using MSAs (Multiple Sequence Alignment) created by MAFFT. It was observed that the pattern of occurrence of TNP-CTDs (Fig. 5) followed the order of evolution: Cnidaria, followed by Arthopoda. THAP9-CTDs (Fig. 6) also follow an evolutionary hierarchy of Arthopoda to Chordata wherein organisms from Amphibia, Reptilia, Aves and Mammalia clustered together thus illustrating higher conservation among member organisms of the same class. However, few organisms from class Chordata (from Pisces: Myxini, Cladistia) were interspersed within Arthropoda.

**Figure 5:**
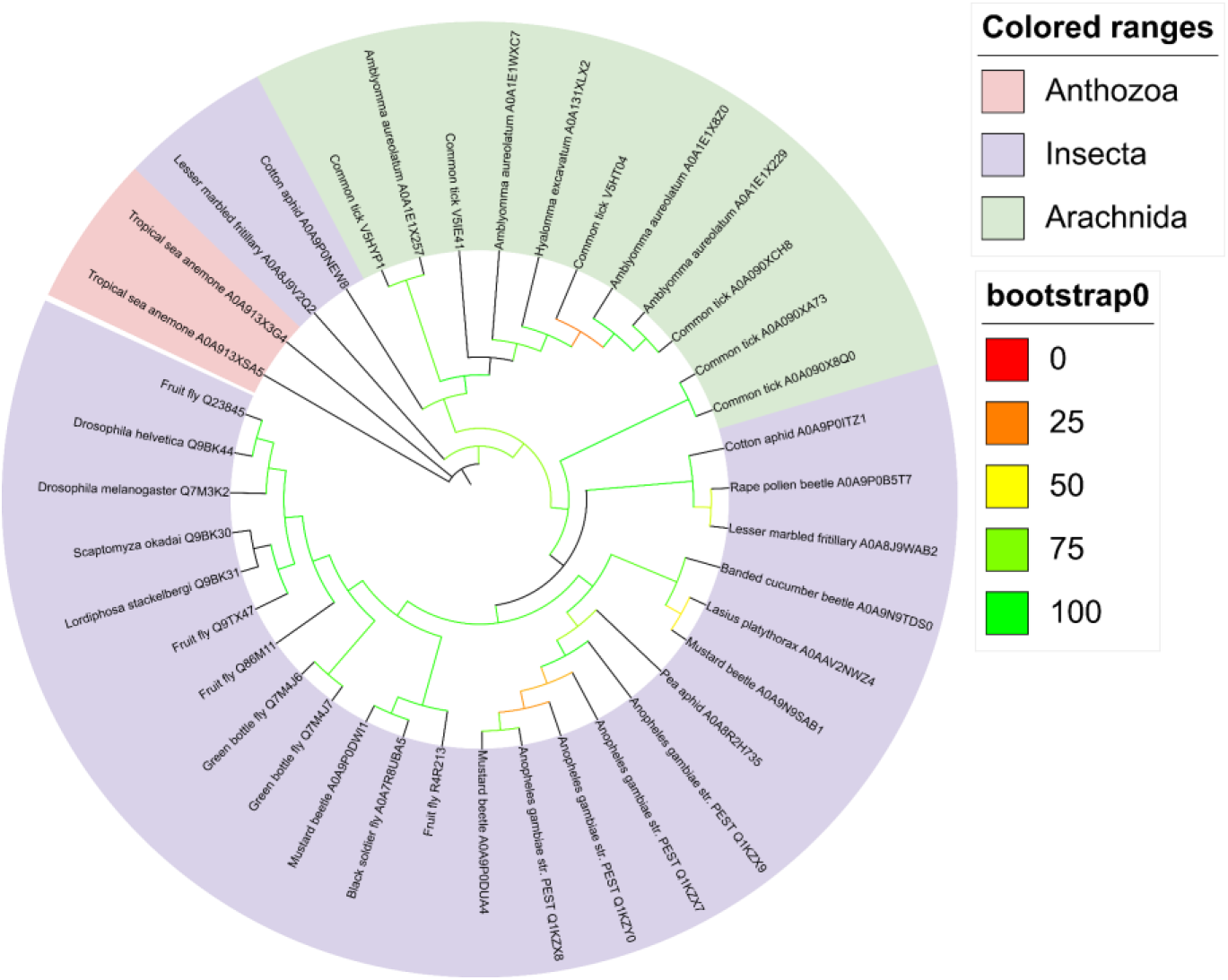
Phylogenetic analysis of TNP-CTD homologs after CD-HIT: The sequences were aligned using MAFFT and the tree was generated using IQ TREE

**Figure 6:**
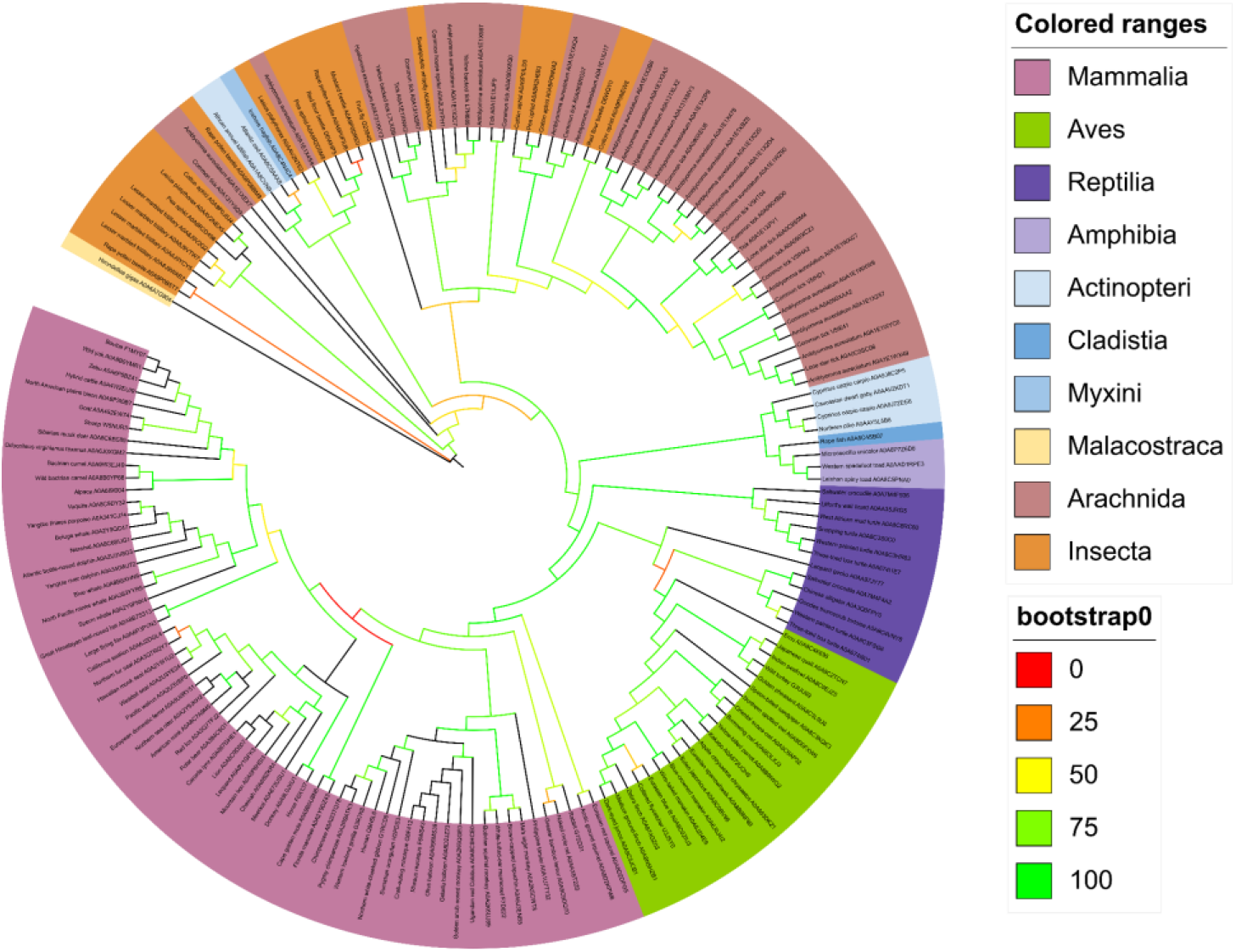
Phylogenetic analysis of THAP9-CTD homologs after CD-HIT: The sequences were aligned using MAFFT and the tree was generated using IQ TREE

### Clustering of TNP-CTD and THAP9-CTD datasets

The homologs of TNP-CTDs and THAP9-CTDs were then individually clustered (using DIAMOND, clustering was performed at 40% sequence similarity) to club organisms possessing similar domains in one group. In a cluster, the centroid is the representative sequence or structure that most accurately captures the characteristics of every other member. This representative sequence was extracted for each group, thereby reducing redundancy and the volume of the datasets. This led to the identification of 38 distinct clusters from the 39 TNP-CTD sequences (Suppl. Table 1) and 75 distinct clusters (Suppl. Table 2) from the 173 THAP9-CTD sequences. It is to be noted that while the clustering of the THAP9-CTDs led to significant decrease in dataset volume (173 to 75), the TNP-CTDs failed to form clusters containing more than one member (Except one variant of fruitfly (29) which formed a centroid with 2 members). This suggests that the TNP-CTD homologs were more divergent. This is also illustrated in the phylogenetic tree (Suppl. Fig 4) which demonstrates that TNP-CTD homologs of Arthropoda tend to form multiple branches rather than large clusters.

On the other hand, for THAP9-CTDs (Suppl. Table 2, Suppl. Fig 5), all mammals except Bolivian squirrel monkey, were clustered into a single entry (centroid as Northern sea otter; 64 members) and all birds were clustered into a single entry (centroid as Zebra finch; 21 members). This suggested that there was strong class-specific sequence conservation of THAP9-CTD within mammals and aves. However, sequence conservation was much less for reptiles [5 clusters: Goodes thornscrub tortoise (1 member), Western painted turtle cluster (4 members), Leopard gecko (1 member), Lilford’s wall lizard (1 member) and Saltwater crocodile (5 members)], amphibians [2 clusters: Western spadefoot toad (2 members), Microcaecilia unicolor (1 member)], fishes [5 clusters] and Arthropods.

The structures of the representative sequences obtained after DIAMOND clustering were then predicted using AlphaFold. The majority of the representatives, for both TNP-CTD and THAP9-CTD, were observed to possess a central core of α helices (predicted with high confidence, marked in blue), which is conserved across homologs. The α helices, varying mostly from 1-3 in number (Suppl. Fig 6 and 7), are joined by weakly conserved loops (predicted with low confidence). No class-specific pattern was observed with respect to the number of helical regions present per homolog.

### Analysis of the regions predicted with high confidence in the CTD homologs

The regions within the AlphaFold-predicted structures of each representative TNP-CTD or THAP9-CTD homolog that were predicted with “very high confidence” (deep blue) and “high confidence” (sky blue) were then further analysed. The corresponding amino acid sequence was retrieved by manual inspection and marked (capital letters for “very high confidence regions”, lower case letters for “high confidence regions”). As observed in Suppl. Fig 6 and 7, the TNP-CTD clusters had three or less “high and very high confidence regions”. These blocks of confidence may not occur simultaneously in an organism but can also exist alone or in combination with other blocks.

### Multiple sequence alignment of TNP-CTDs and THAP9-CTDs illustrate conserved amino acid residues across all classes

Multiple sequence alignment (MSA) of the amino acid sequences of the cluster representatives (38 for TNP-CTDs, 75 for THAP9-CTDs) was performed and the identified high and very high confidence regions were mapped. Interestingly, it was observed that the helical regions within the AlphaFold-predicted structures that were predicted with “very high confidence” overlapped with with three conserved blocks of hydrophobic (blue) amino acid sequences (Regions 1, 2, 3) in both TNP-CTDs (Fig.7) as well as THAP9-CTDs (Fig.8). These “high confidence”regions were divided into parts “a” and “b”, where “a” corresponded to the first two conserved blocks (Regions 1, 2) which appeared almost simultaneously, and were sometimes separated by less confident loops while “b” corresponded to conserved Region 3. Interestingly, this pattern of conservation was also observed when MSA (Suppl. Fig.1, 2) was performed with all the homologs (i.e. before clustering).

**Figure 7:**
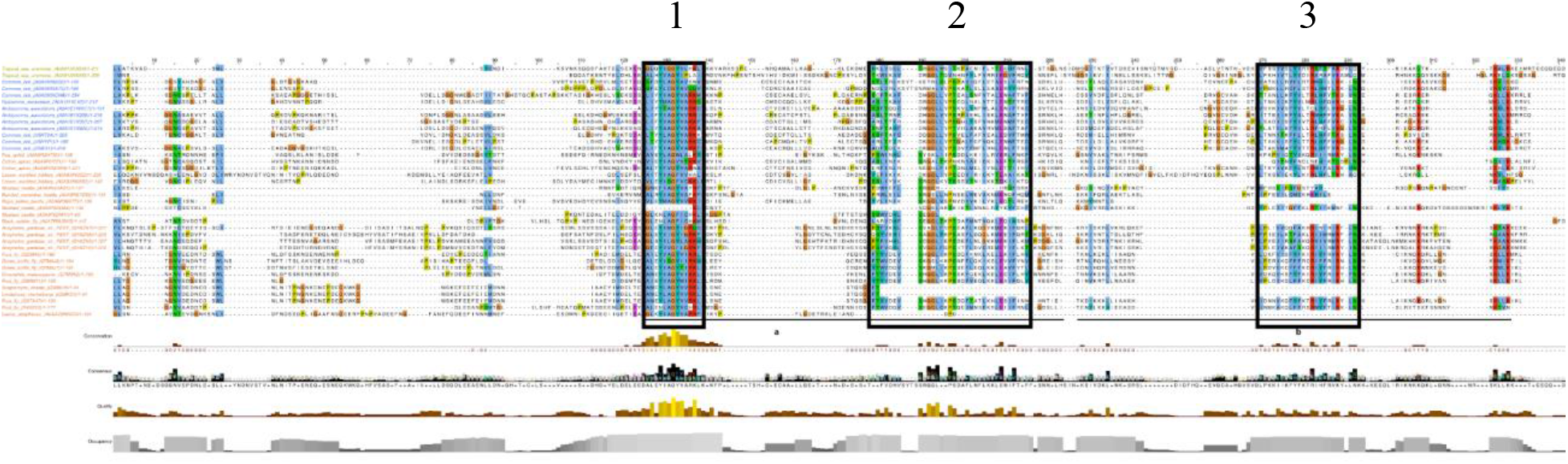
MSA of TNP-CTD centroid clusters with conserved blocks 1,2,3 (black-outlined boxes) coincides with AlphaFold’s high and very high confidence regions. The names of the species have been colored as per their classes: Anthozoa (yellow), Arachnida (royal blue), Insecta (burnt orange).

**Figure 8:**
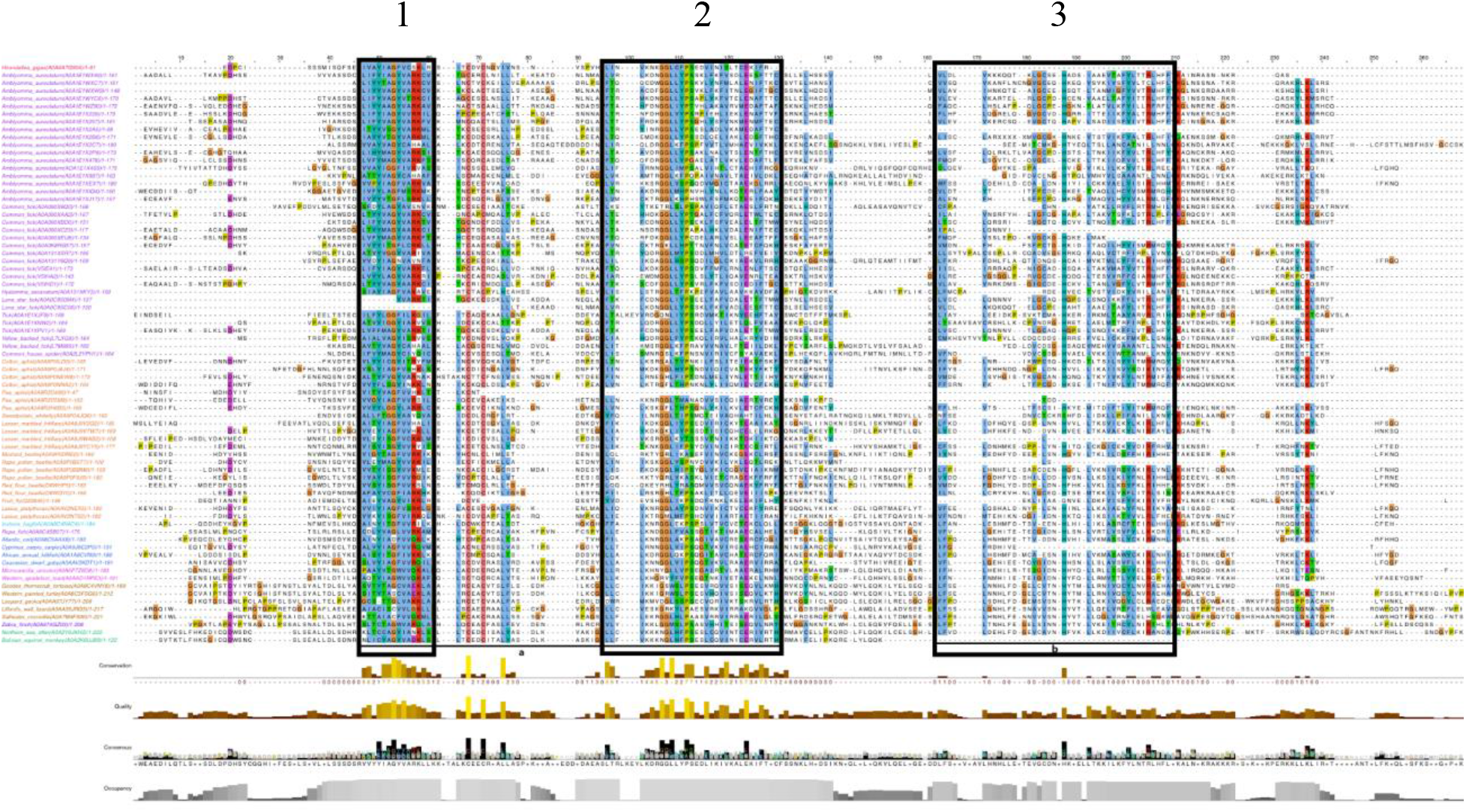
MSA of THAP9-CTD centroid clusters with conserved blocks 1,2,3 (black-outlined boxes) coincides with AlphaFold’s high and very high confidence regions. The names of the species have been colored as per their classes: Malacostraca (red), Arachnida (bright purple), Insecta (burnt orange), Myxini (teal), Clastidia (violet), Actinopteri (blue), Amphibia(pink), Reptiles (orange), Aves (purple), Mammals (green). The panel below shows sequence conservation, alignment quality, consensus sequence, and residue occupancy across the MSA.

Further, detailed observation of the three evolutionarily conserved regions illustrated that they are characterized by distinct combinations of hydrophobic, aromatic and charged residues. Region 1 is enriched in small hydrophobic and aromatic residues, Region 2 contains a mixed hydrophobic/basic composition with glycine-and proline-associated boundaries, whereas Region 3 represents the most extensive conserved hydrophobic/charged block, containing repeated leucine, isoleucine, valine and phenylalanine residues interspersed with conserved acidic and basic residues. These features are consistent with a helical structural core in which hydrophobic residues contribute to intradomain packing while charged residues help to stabilize the protein fold through solvent exposure or electrostatic interactions (Zhou and Pang 2018). The conserved consensus motif corresponding to each helical region in TNP-CTDs (Suppl Fig 1), are ALEYIAGYVARKL (Region 1), FTFVDHVSRGGLIKPSDAFLNFLKKLENIFTXF (Region 2) and LPKKIIKFYFKTRIHFRVKYLN (Region 3).The conserved consensus motif corresponding to each helical region in THAP9-CTDs (Suppl Fig 2), are LTYCAGYVANKL (Region 1), LLCVKKKGGLHFPSESLCRVINICERVVRTH (Region 2) and LFVDLDEHLFDGEVCAINHFVKLLKDIIICFLKIRAKDV (Region 3).

To investigate if there is any conservation between the THAP9-CTDs and TNP-CTDs, a combined MSA was also performed with both datasets together (Suppl. Fig. 3). Interestingly, it was observed that the conserved regions in TNP-CTDs perfectly aligned with the conserved helices of THAP9-CTD (Block 1,2,3 in Suppl. Fig 3) thus illustrating an overlap of secondary structural elements in the CTDs of DmTNP and THAP9 homologs.The horizontal block in Suppl. Fig 3 represents the 12 organisms (all arthropods) in which the identified protein homolog has a CTD that is homologous to the CTDs in both DmTNP and hTHAP9 (is retrieved from Uniprot using both “DmTNP-CTD” and “hTHAP9-CTD” as query). It is tempting to suggest that these 12 arthropods could form the junction for the evolutionary divergence between TNP-CTDs and THAP9-CTDs. The conserved consensus motif corresponding to each helical region for both TNP-CTDs and THAP9-CTDs (Suppl Fig 3), are LTYIAGYVARKL (Region 1), LLCVKKKGGLHFPSESLCRVINICERVVRTH (Region 2) and ILCELSGHIYLFVDLDEHLFDGEVCAINHFVKLLKDIIICFLKIRA (Region 3).

It was noted that THAP9-CTDs have greater sequence conservation among birds and mammals and lesser sequence conservation among reptiles, amphibians, fishes, annelids, cnidaria, arthropods (Fig. 8). The extended C-terminal end of THAP9-CTD is approximately 25 amino acid residues and is highly-conserved among mammals (Suppl. Fig 2). Moreover, the N-terminal end of THAP9-CTDs appear to have class-specific conservation (in Reptiles, Aves and Mammals) such that they are more similar for members of the same class as compared to members of other classes. For example, the N-terminal end of THAP9-CTD in few members of Reptilia have a conserved motif: Gxxxxx(S/T)xDYxYxxG, Aves have A(R/P/H)GR(T/M)(L/P/V)(A/P/T)x[E/K/G/Q]YPx(C/R)AG while mammalis have a hydrophobic motif: S(V/I)(I/V/T)K(T/S)(L/V)FH(K/E)E(D/G).

### Multiple Structural Alignment (MSTA) of Clusters indicate the homologs of THAP9-CTDs are more closely related than DmTNP-CTDs

To investigate if there was structural conservation amongst the individual cluster members, Multiple Structural Alignment (MSTA) was performed for both the CTDs using MUSTANG. RMSD (Root Mean Square Deviation) tables (Konagurthu et al. 2006) were generated and heatmaps for each cluster were prepared with the RMSD values wherein an RMSD of 4 Å or below was considered as high similarity (Fig. 9). The superimposed structures from MUSTANG are shown in Fig. 9a and c for TNP-CTDs and THAP9-CTDs respectively. Both sets of homologs found a common core to superimpose suggesting that the structure is similar to each other. As per the key, the increase in darkness of blue colour on the heatmap corresponds to how closely related the representative structures are. The centroid clusters in both the heatmaps are numbered according to Suppl. Table 1 and 2.

**Figure 9:**
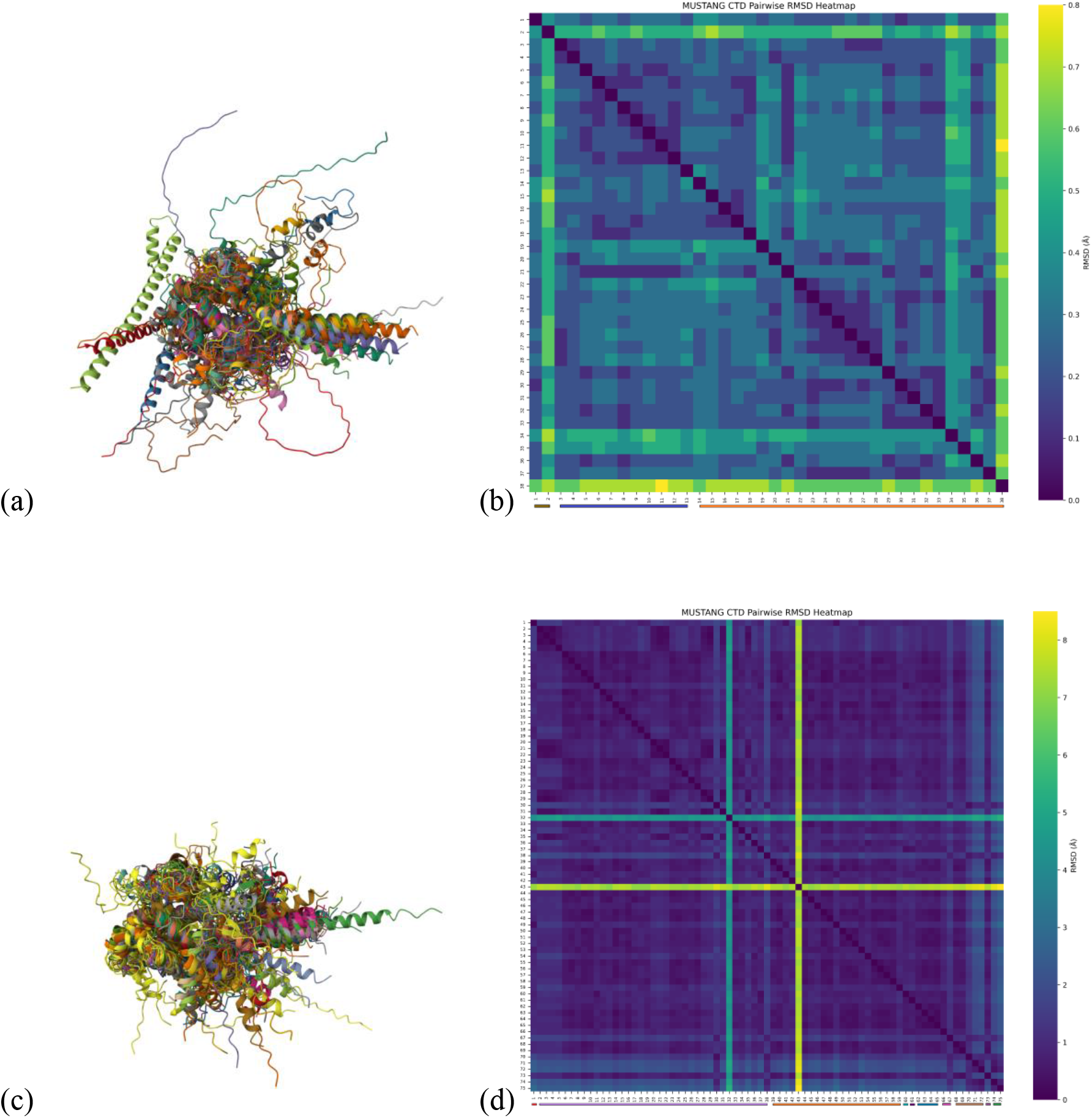
Superimposed structures of all the centroid sequences of (a) TNP-CTDs and (c) THAP9-CTDs obtained from MUSTANG. Heatmap of all the (b) TNP-CTD and (d) THAP9-CTD clusters, where the right panel shows RMSD values. indicating more similarities within members of the same phylum (blue) and lesser similarities with members of different phylum (yellow). The organism numbers indicated in the axes of (b) and (d) can be referred from Suppl. Tables 1 and 2 respectively. The species nos. have been colored as per their classes: Anthozoa (yellow), Arachnida (royal blue), Insecta (burnt orange) - in (b) and Malacostraca (red), Arachnida (bright purple), Insecta (burnt orange), Myxini (teal), Clastidia (violet), Actinopteri (=blue), Amphibia(pink), Reptiles (orange), Aves (purple), Mammals (green) - in (d).

It is observed that amongst the TNP-CTD homologs (Fig. 9b), all the members (except species no. 1 and 2) belong to the phylum Arthropoda (Insecta and Arachnida) and possess highly similar CTD structures with RMSD values less than 1Å. The colour bar below the heatmap shows the class of the organisms. Amongst the THAP9-CTD homologs, most vertebrates - from amphibians, reptiles to mammals, (Fig 10d) possess a conserved CTD structure (in heatmap, separate dark blue RMSD block from organism numbers 67 to 75) except for aves. This indicates that the CTD of most vertebrates are similar to each other. Aves have slightly lesser similarity with mammals and reptiles but more similarity among other members, hence a distinction from the dark blue RMSD block (indicated by light blue). Arthropods (Insecta and Archnida), too, show dense RMSD among different species of the same class, and also with members to pisces and amphibia, indicating strong conserved structure, but higher RMSDs when compared with other vertebrates (dark blue block from organism no 2 to 66).

**Fig 10:**
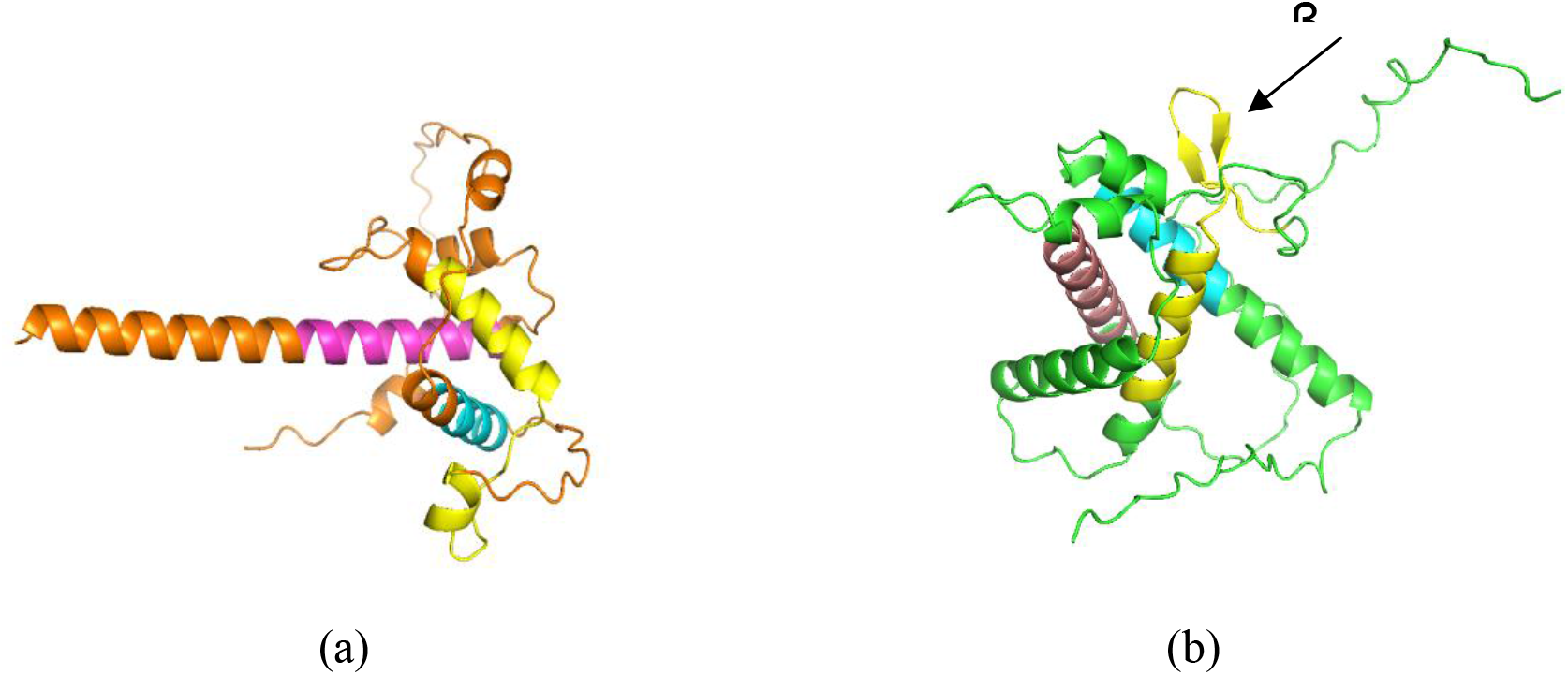
Alphafold predicted structures of CTDs of (a) DmTNP and (b) hTHAP9. The regions marked in cyan, yellow and pink (a and b) correspond to conserved hydrophobic regions 1, 2 and 3 from Suppl. Fig 1 and 2, and are predicted to arrange into α helices, forming a putative structural core.

In Fig. 9d, Lone star tick and pea aphid show higher RMSD as compared to other organisms (indicated by the presence of teal and yellow lines in the heatmap respectively). Lone star tick (organism no. 32), despite having three α helices, failed to align with other sequences. This could be due to the difference in number of residues contributing to the formation of each α helix rather than the total protein length. Pea aphid (organism no. 43), on the other hand, has a very small 47-residue CTD which fails to align with the common core and hence shows high RMSD value.

### Structural analysis of hTHAP9-CTD and DmTNP-CTD

Investigation of the AlphaFold predicted structures of DmTNP and THAP9-CTDs demonstrated that hTHAP9-CTD was longer (by 38 amino acids) than DmTNP-CTD (Fig. 10). Here too, the regions with high confidence scores (marked in cyan, yellow and pink) (a and b) correspond to conserved hydrophobic regions 1, 2 and 3 and are predicted to arrange into α helices, forming a putative structural core while the interconnecting loops and β-sheets had low confidence scores. The two ends of the DmTNP-CTD are not well-modelled in the DmTNP cryo-EM structure possibly due to the higher probability of disorderliness in these regions (Ghanim et al. 2019).

### Both the terminal regions of CTDs of DmTNP and THAP9 show disorderliness

Although structural predictions of CTDs of both DmTNP and hTHAP9 indicate the presence of conserved folded and coiled-coil regions, intrinsically disordered regions can also exist simultaneously with structured elements and often play essential roles in protein–protein interactions, regulatory processes, and post-translational modifications (Wright and Dyson 2015). IUPred analysis demonstrated that overall the DmTNP-CTD was more disordered than hTHAP9-CTD (Fig. 11a). Further, the N-terminal and C-terminal tails of each CTD had more disorder (above the disorder threshold) but the magnitude of disorder was higher for DmTNP than THAP9. For example, hTHAP9-CTD has disorder between residues 1-22 and 205-210 whereas DmTNP has disorder between residues 11-43 and 97-100 as well as the end. Moreover, the disordered architecture was conserved (Fig. 11b and c) across the TNP-CTD and THAP9-CTD homologs, where the transition between order and disorder occur synchronously. The ordered regions in Fig. 11 b and c correspond to the conserved helical regions in Suppl. Fig 1 and 2 for TNP-CTD and THAP9-CTD homologs whereas the disordered regions are in accordance with the non-conserved regions.

**Figure 11:**
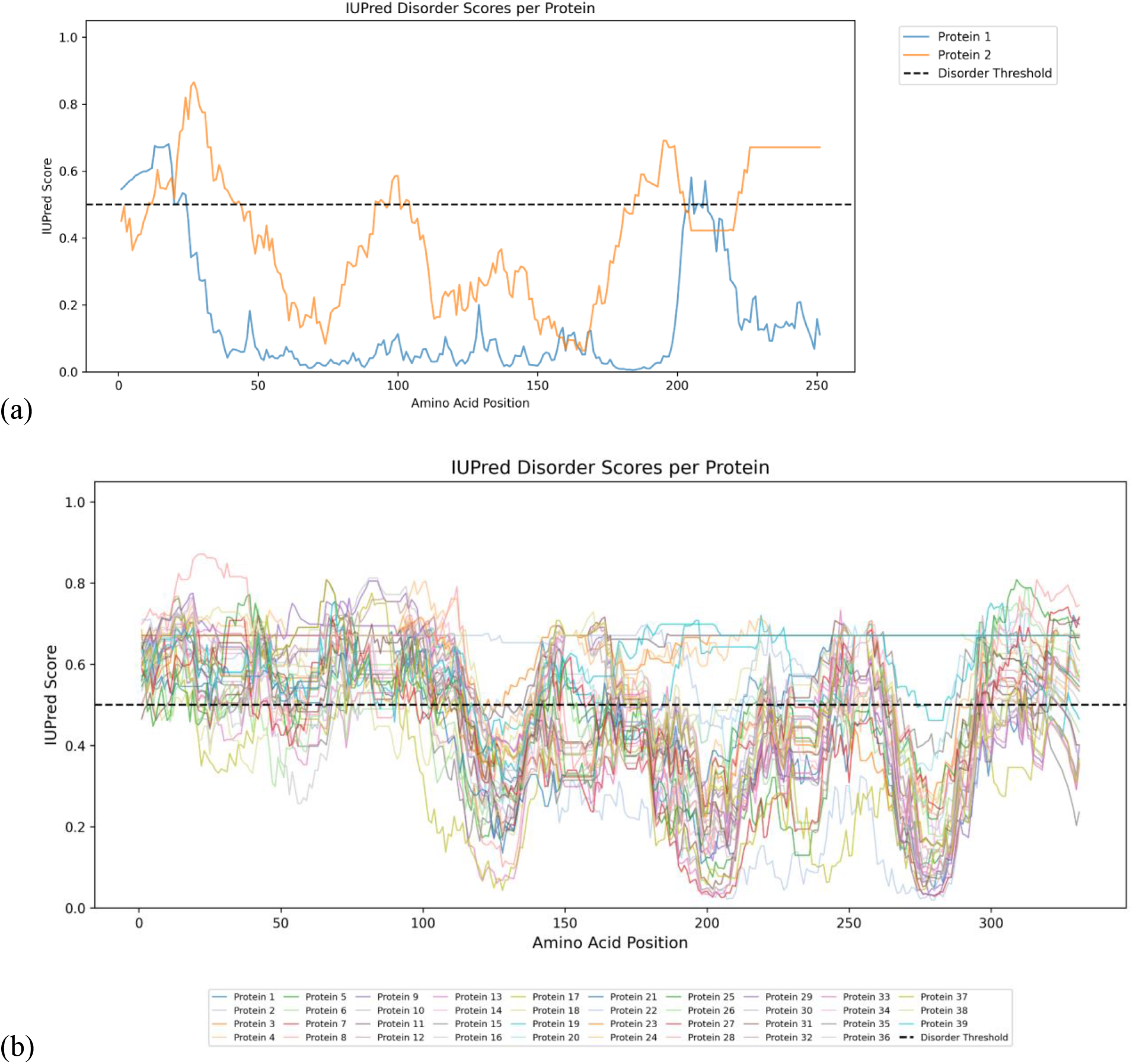
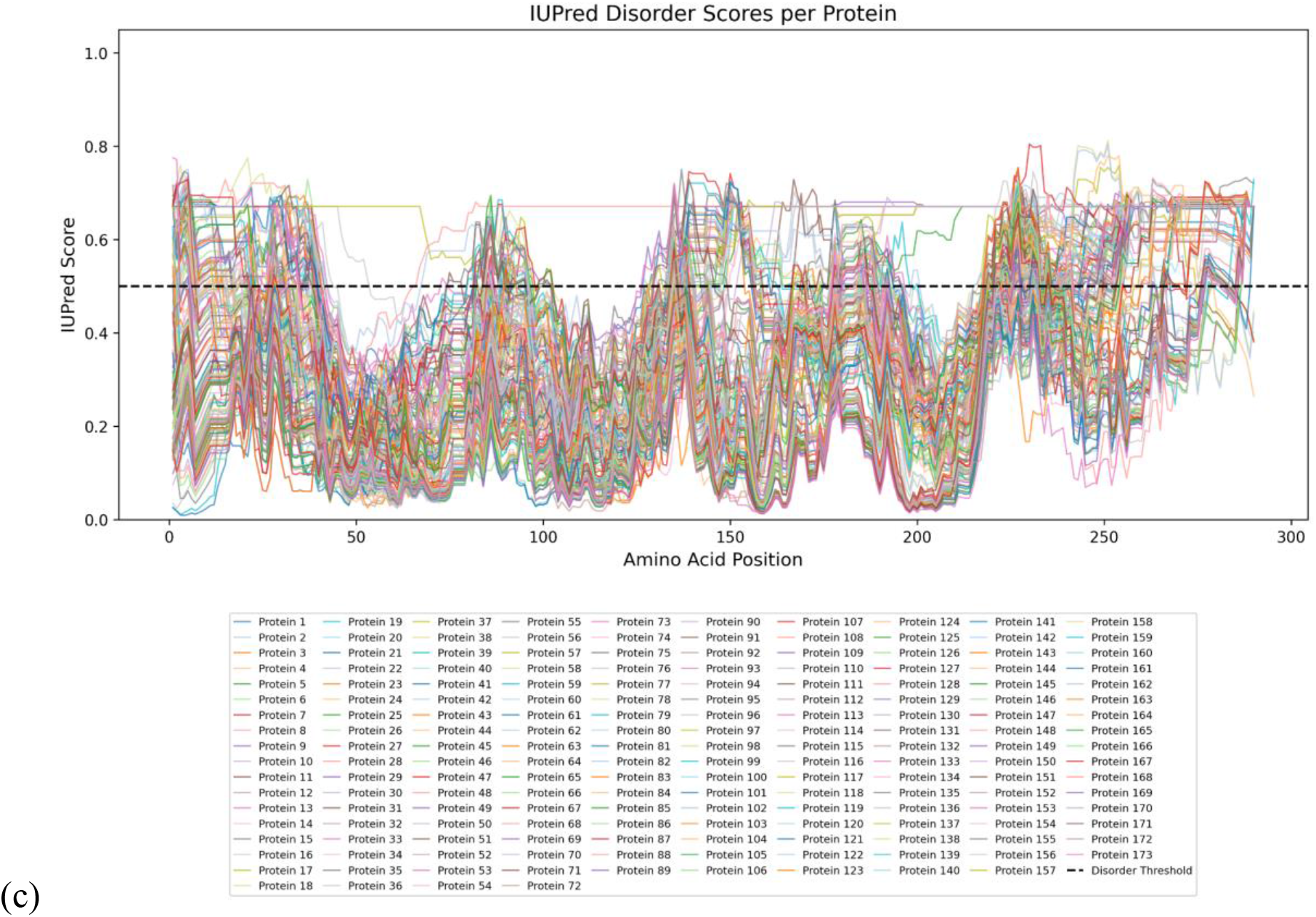
IUPred analysis demonstrating intrinsic disorder landscape in (a) DmTNP (Protein 2) and THAP9 (Protein 1) (b)TNP-CTD homologs (c) THAP9-CTD homologs

**Figure 12:**
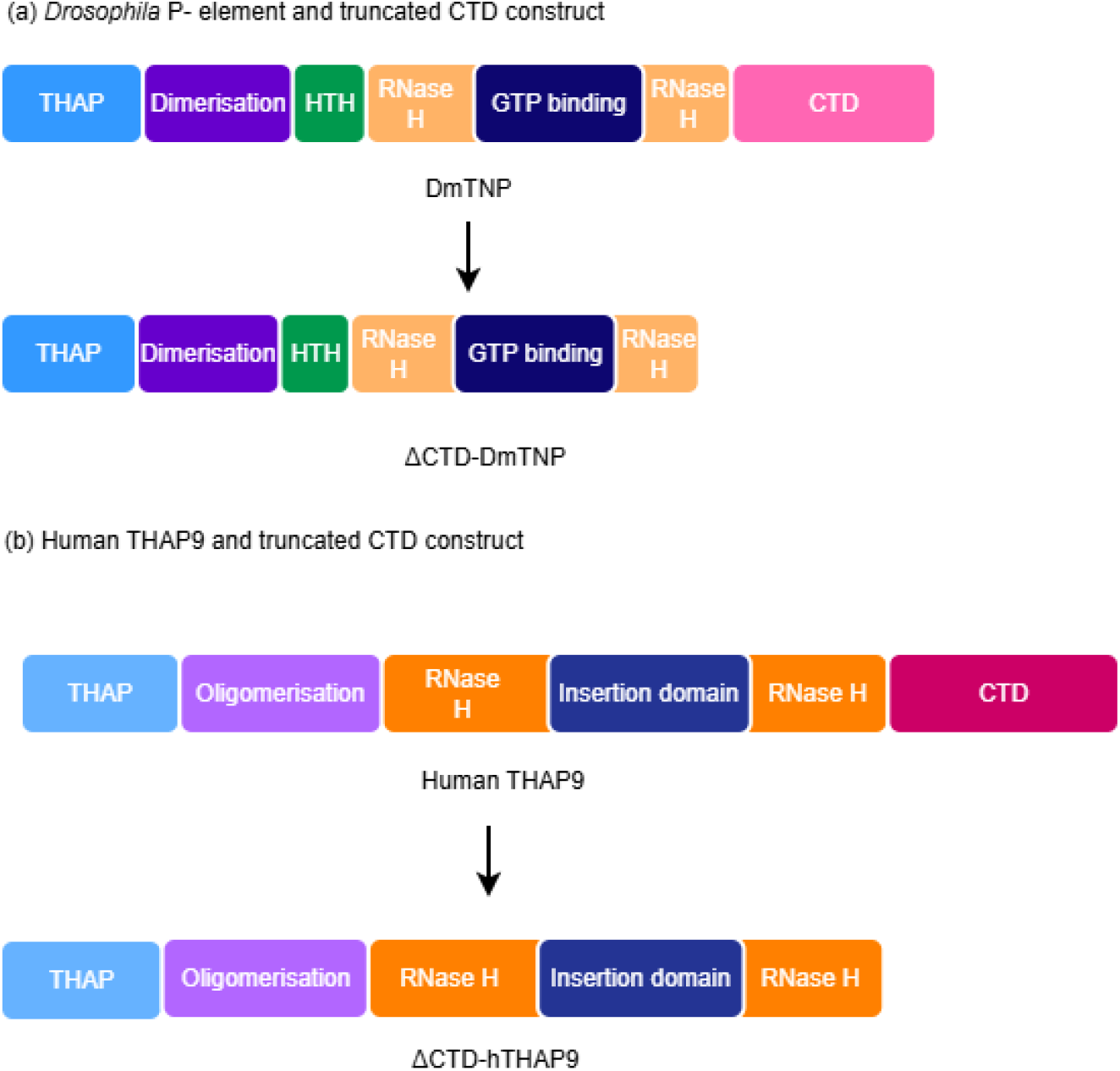
Schematic representation of the CTD mutant constructs of DmTNP(a) and hTHAP9(b) used for DNA excision and integration assays

### Truncation of C-terminal domains of hTHAP9 and DmTNP

Thus, our extensive phylogenetic analyses on the CTDs of DmTNP and hTHAP9, suggested that these domains contained novel folds with strongly conserved helical regions. However, the biochemical role of this domain in both DmTNP and hTHAP was still unknown. To investigate the roles of the CTDs of hTHAP9 and DmTNP, truncated mutants were created in which the CTD was deleted.

For DmTNP, C-terminal domain mutant was truncated from amino acids 567-751. This boundary was determined from the solved cryo-EM structure (Ghanim et al. 2019). For hTHAP9, the C-terminal domain mutant was truncated from amino acids 681-903. Here, the C-terminal boundary was determined based on the domain boundary obtained from AlphaFold.

### The CTD of hTHAP9 is not required for DNA cleavage

To investigate the catalytic activity of ΔCTD-hTHAP9 and ΔCTD-DmTNP mutants, plasmid-based DNA excision assays were performed. A reporter plasmid (pISP-2/Km, having 0.6 Kb insertion of P-element DNA flanked by DmTNP TIRs) was co-transfected with a plasmid encoding the transposase. The DNA excision activity was confirmed by the ability of the transposase (Wild type as well as mutants) encoded by a transfected plasmid to excise the P-element DNA after binding the flanking TIRs. Upon successful excision, the reporter plasmid would be repaired by the cellular repair machinery. Hence, after isolation from the cells, the plasmid would yield a shorter band (∼200 bp) upon PCR amplification across the TIR region as compared to the ∼750bp band that is obtained if P-element excision has not occurred.

As observed in Fig.13, both ΔCTD-hTHAP9 and ΔCTD-DmTNP mutants gave rise to similar PCR amplification patterns of the reporter plasmid (200 bp) like the wildtype full length hTHAP9 and DmTNP. This indicates that both hTHAP9 and DmTNP can successfully recognise and excise the P-element DNA (Fig. 13) in the absence of CTD. It is interesting to note that although DmTNP-CTD has been shown to contact DNA in the strand transfer complex (Ghanim 2019), the domain is dispensable with respect to its ability to cleave DNA.

**Figure 13:**
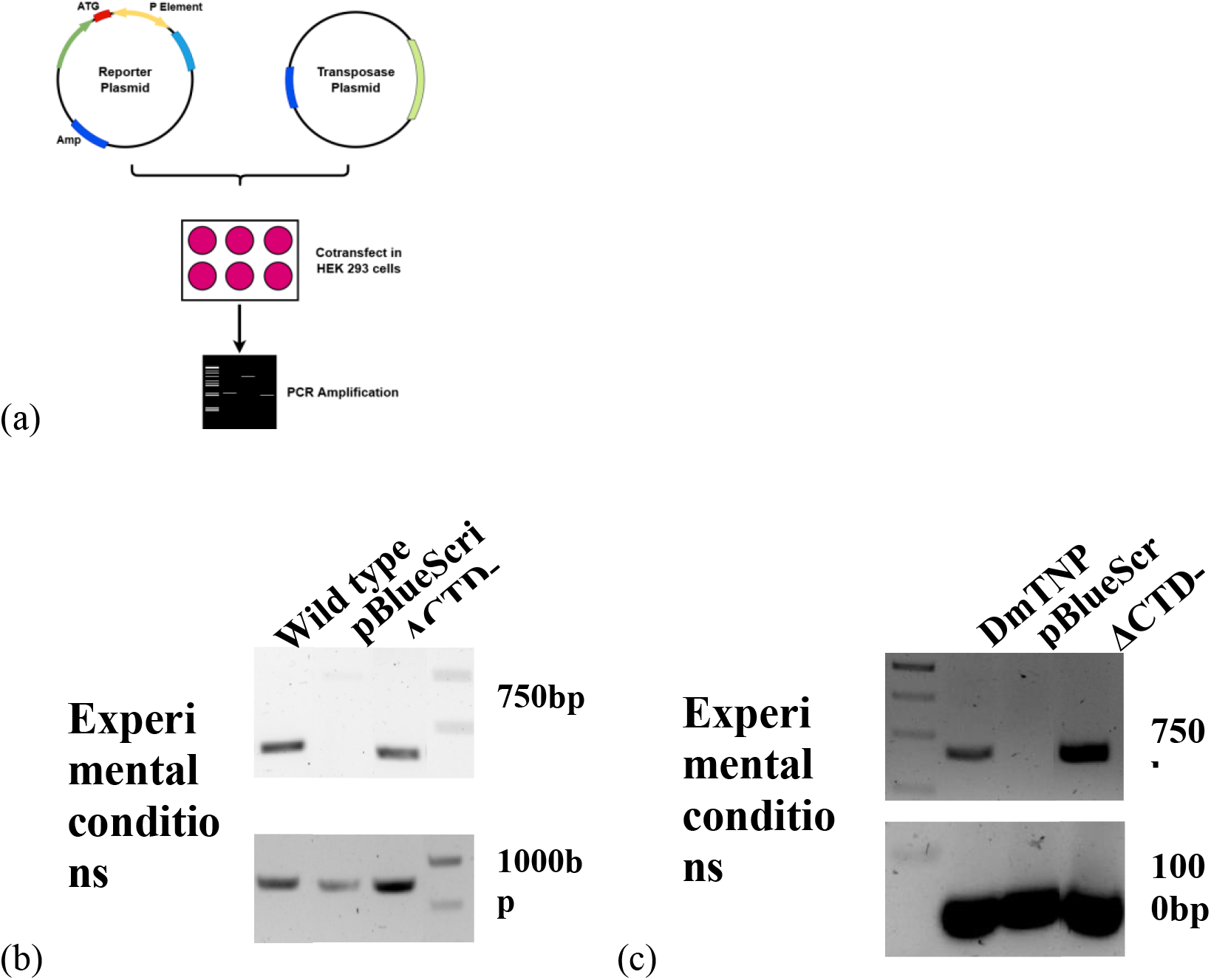
(a) Schematic of DNA excision assay. DNA excision ability of hTHAP9 (b) and DmTNP (c) and their truncation mutants with Ampicillin resistance gene as template control.

### The CTD of hTHAP9 regulates its ability to integrate DNA

After confirming that CTD deletion did not disrupt hTHAP9’s ability to recognize and cleave P-element DNA, we next investigated whether the domain was required for DNA integration. The ability of the transposase source (wild type or mutants) to excise and then integrate the G418 resistance cassette flanked by P-element TIRs was assessed by screening and staining G418-resistant colonies (described in the Methods section). The relative DNA integration activity (Suppl. Table 3, Sharma et al. 2021) was then calculated.

Interestingly, it was observed that ΔCTD-hTHAP9 retained the ability to integrate DNA. Moreover, its DNA integration activity was enhanced by ∼60% (Fig. 14b) when compared to wildtype. On the other hand, deletion of the CTD in DmTNP (ΔCTD-DmTNP) did not affect its ability to integrate DNA; its integration activity was at par with the wild type DmTNP.

**Figure 14:**
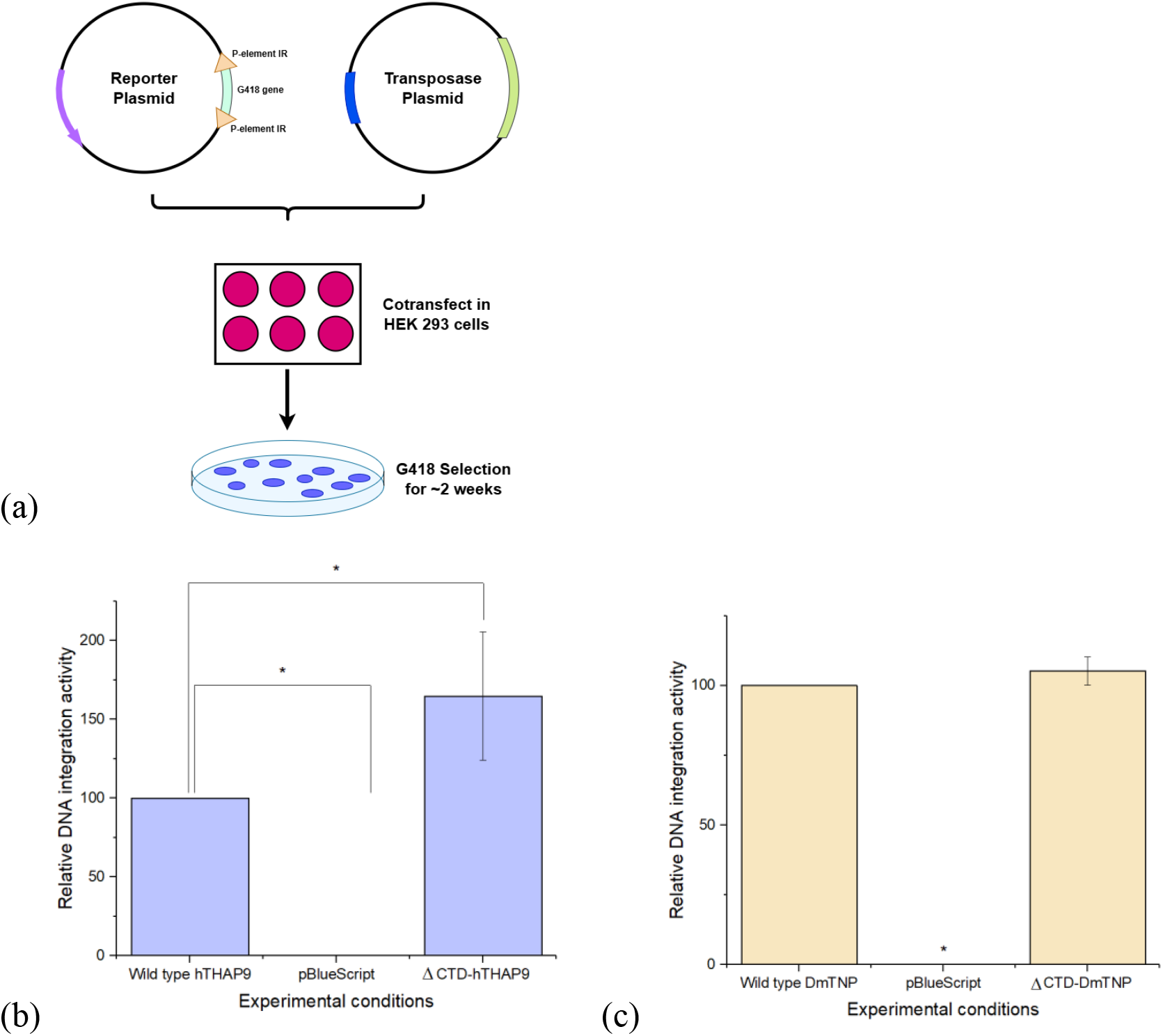
(a) Schematic of DNA integration assay. Relative DNA integration activity of (b) hTHAP9 and (c) DmTNP and their CTD-truncation mutants; pBluescript=negative control.

Thus, the results for DNA integration assay are in correspondence with that of the DNA excision assay, suggesting that both the truncated mutants of THAP9 and DmTNP retain the ability to mobilise P-element DNA (Fig.14). This suggests that the C-terminal domains in both hTHAP9 and DmTNP, are not required to excise and integrate DNA. Moreover, the hTHAP9-CTD may be negatively regulating DNA integration activity since its absence enhances hTHAP9’s ability to integrate DNA.

This study employed a combination of bioinformatic and biochemical approaches to investigate the CTDs of hTHAP9 and DmTNP. The bioinformatic analysis demonstrated that both the domains adopt distinct, hydrophobic helical regions that are conserved across diverse groups of organisms having the homologs of the domain. The biochemical assays demonstrate that the CTD of hTHAP9, and not DmTNP, significantly inhibits its ability to integrate DNA and thus may play a regulatory role.

## Discussion

The terminal domains of a protein are often crucial for its functioning, cellular localisation as well as its stability. The N-terminal domain serves as a site for post translational modification, harbouring sites for cellular location as well as degradation of proteins (Walling 2006). The C-terminal domain also plays a role in maintaining protein quality check, its stability as well as degradation (Chu et al. 2026). Hydrophobic residues within helical regions in C-terminal domains of proteins have been observed to maintain the overall three dimensional structure of the protein as well as its functionality. For example, in Apolipoprotein A-1, the C-terminal helix bundle domain interacts with the N-terminal helix bundle domain maintaining the conformation of the protein (Lyssenko et al. 2012). Here we investigate the uncharacterised C-terminal domains (CTD) of the *Drosophila* P-element transposase (DmTNP) and its homolog, hTHAP9.

We observe that the CTDs of DmTNP and hTHAP9 do not have high sequence identity. However, they both have similar secondary structures comprising three hydrophobic alpha-helical regions (Fig. 7, 8) that are conserved in homologs from diverse organisms. This illustrates that their individual structures may be more conserved than their underlying amino acid sequence, as observed in other proteins (Rajendran and Jothi 2018) including various transposase-derived proteins, where the fold of the protein is more preserved than the sequence (Abrusán et al. 2013). We investigated if any of these helical regions can form coiled coil structures in DmTNP and hTHAP9, which are often responsible for protein-protein interactions. However, DeepCoil2.0 (Ludwiczak et al. 2019) predicted that the probability of the CTDs forming coiled coil structures is very low (data not shown).

Interestingly, as per the Interpro database and previous work (Rashmi et al. 2024), both the CTDs of DmTNP and THAP9 are novel folds which do not exist in any other protein apart from homologs of DmTNP and THAP9 respectively. IUPred analysis (Fig. 11) illustrates that the C-terminal tails of these CTDs are disordered and hence could play regulatory functions. For example, they could serve as sites for post translational modifications (as seen in cAMP regulated CREB transcription factor which is activated by the phosphorylation of Ser133 residue in its intrinsically disordered kinase inducible activated domain), promiscuous motifs for protein-protein interactions (as seen in Adenoviral oncoprotein early region 1 (E1A) in which 3 different intrinsically disordered regions recruit different proteins), or maintain structure (Wright and Dyson 2015).

The cryo-EM structure of the DmTNP paired-end strand transfer complex (PDB ID: 6PE2) illustrates that the DmTNP-CTD is involved in DNA-protein interactions with the strand transfer DNA (Ghanim et al. 2019) via residues K637 and L661 which align with K725 and L761 of hTHAP9. Interestingly these residues are also conserved across all TNP-CTD and THAP9-CTD homologs and lie in Region 3 (Fig 8). However, our analysis of CTD-truncation mutants demonstrates that the deletion of DmTNP-CTD has negligible impact on its DNA excising and integrating ability. This suggests that the DmTNP-CTD has an auxiliary but not essential role in DNA binding, which can be compensated by other domains.

On the other hand, deletion of CTD in hTHAP9 leads to an increase in its DNA integration efficiency, thus implicating that this domain may function as a negative regulatory domain. Based on these observations, it is tempting to speculate that the acquisition of a novel CTD by THAP9 has helped to regulate its enzymatic activity and prevent uncontrolled transposition. The hTHAP9-CTD may thus be an auto inhibitory module, which controls hTHAP9 or putative interaction partners that mediate the DNA cut-and-paste activity. This behaviour is also in accordance with the hypothesis that THAP9 is a transposon derived gene which has been domesticated, but retains the catalytic activity with several layers of control and modulation.

The THAP9-CTD has a novel fold that is not observed in any other protein and is distinct from the CTD of the DmTNP. Thus, it is hypothesised that the domain may have evolved by the process of exonisation (e.g., *Alu* retrotransposable elements acquire new functional exons via genetic integration, substitution and deletion during the course of evolution (Sorek 2007) or domain accretion (e.g., fusion of a domain or peptide as seen in FokI which has acquired a C-terminal catalytic domain, homologous to EcoRV, thereby recognising the EcoRV recognition sequence 5’-GGATG-3’ (Bashton and Chothia 2007).

This study sheds light on the evolution of the CTDs of DmTNP and hTHAP9 and its homologs. The domain is structurally conserved but sequence-divergent and has different roles in DmTNP and hTHAP9: it supports DNA binding in DmTNP whereas in hTHAP9, it acts as a negative regulator. In the future, it will be intriguing to understand the biological role of the hTHAP9-CTD as well as its potential nucleic acid and protein binding partners.

